# A generic decision-making ability predicts psychopathology in adolescents and young adults and is reflected in distinct brain connectivity patterns

**DOI:** 10.1101/2020.08.20.259697

**Authors:** Michael Moutoussis, Benjamín Garzón, Sharon Neufeld, Dominik R. Bach, Francesco Rigoli, NSPN Consortium, Marc Guitart-Masip, Raymond J. Dolan

**Affiliations:** Wellcome Centre for Human Neuroimaging, University College London, London WC1N 3BG, United Kingdom; Max Planck University College London Centre for Computational Psychiatry and Ageing Research, London WC1B 5EH, United Kingdom and 14195 Berlin, Germany; Aging Research Centre, Karolinska Institute, Stockholm, Sweden; Department of Psychiatry, University of Cambridge, Cambridge CB2 0SZ, United Kingdom; Computational Psychiatry Research; Department of Psychiatry, Psychotherapy, and Psychosomatics; Psychiatric Hospital; University of Zurich, 8032 Zurich, Switzerland; Department of Psychology, City University, London, United Kingdom; The list of the NSPN Consortium members can be found in part F of the Supplementary material

## Abstract

Decision-making underpins many important facets of our lives. Here, we assessed if a general ability factor underpins decision-making abilities. Using factor analysis of 32 decision-making measures in 830 adolescents and young adults, we identified a common factor we refer to as ‘decision acuity’ that was distinct from IQ and reflected advantageous decision-making abilities. Decision acuity decreased with low general social functioning and aberrant thinking. Crucially, decision acuity and IQ had dissociable neural signatures in terms of resting-state functional connectivity involving specific neural networks. Finally, decision acuity was reliable and its relationship with functional connectivity was stable when measured in the same individuals 18 months later. We conclude that our behavioural and brain data demonstrate a new cognitive construct encapsulating ability to perform decision-making across distinct domains, and that the expression of this construct may be important for understanding psychopathology.

## Introduction

Decision-making abilities are important for economic performance and social adaptation, and a computational characterization of decision-making processes is likely to advance the understanding of psychiatric disorders (Scholl & Klein-Flügge, 2018). Yet, unlike traditional cognitive constructs such as intelligence, the distribution and covariation of decision-making characteristics in the population is unknown and the reliability of behavioral tasks typically used to measure them has been questioned (Brown et al., 2020; Hedge et al., 2020). Likewise, we know little about the neural underpinnings of decision-making during adolescence and early adulthood, a crucial period for brain maturation (Giedd, 2004; Whitaker et al., 2016). Advancing our understanding here is rendered quite important by the fact that the bulk of psychopathology emerges during adolescence and early adulthood (Paus et al., 2008).

Decision-making involves an interplay of multiple cognitive abilities needed to evaluate available options and settle on a course of action (Kable & Glimcher, 2009; Phelps et al., 2014). Reinforcement-learning has helped characterize the computational and neurobiological processes by which individuals evaluate options (Dayan & Daw, 2008; Sutton & Barto, 1998). This literature is often framed in terms of model-based and model-free evaluations (Daw et al., 2005; Dolan & Dayan, 2013). In the former, the value of different actions is calculated prospectively based on the goals and actions that will lead to these goals. In contrast, the latter involves learning the value of actions by associating them with the value of experienced outcomes.

The relative importance of different evaluation systems is an important individual difference, likely to be trait-like at least in part. Importantly, model based and model free approaches trade off at different levels in different individuals (Eppinger et al., 2017; Kool et al., 2017). Similarly, the impact of Pavlovian heuristics, that is, the propensity to attach value to specific actions by mere association with perceived features of a context, also varies across individuals (de Boer et al., 2019; Guitart-Masip et al., 2012; Moutoussis et al., 2018). Individuals also differ with respect to other factors affecting the evaluation of options, for example, in their aversion to variability of outcomes for an action rather than its mean outcome (Christopoulos et al., 2009; Payzan-LeNestour et al., 2013). Similarly, individuals balance the need to actively collect rewards against the risks of potential dangers in the environment (Bach et al., 2020; Loh et al., 2017; O’Neil et al., 2015). In the temporal domain, individuals balance a need to exploit known choices against uncertainty of exploring unknown ones (Badre, Doll, Long, & Frank, 2012; Sutton & Barto, 1998). Finally, understanding of intentions and emotions of others has a big impact when making decisions in social contexts (King-Casas et al., 2008; Moutoussis, Dolan, & Dayan, 2016).

Although fundamental decision-making characteristics are likely to be largely distinct, we hypothesised that they would also be subject to covariation in the population. In this frame of reference, shared variance along latent dimensions is analogous to the structure of intelligence, where a cornucopia of abilities covaries with latent dimensions such as general and domain-specific intelligence (Van Der Maas et al., 2006). We hypothesised that the main constructs influencing performance across distinct instances of decision-making would include sensitivity to gains and losses, the extent to which model-based approaches dominate choice evaluation, an overall propensity to take risks, and an ability to make good social judgements.

To assess dimensions of decision-making ability, we examined a battery of seven decision-making tasks (table 1), administered to 830 14-24 year olds sampled from a pool of about 2400 young people living in the community in England (Kiddle et al., 2017). We used computational modelling and key descriptive statistics to extract component measures of decision-making (Bach et al., 2014; Fett et al., 2012; Moutoussis, Bentall, El-Deredy, & Dayan, 2011; Moutoussis et al., 2018, 2016; Rigoli et al., 2016; Shahar, Hauser, et al., 2019). We then derived latent cognitive constructs underlying decision-making across tasks, by submitting the component measures to factor analysis (see Methods) and assessed their reliability using the data of 571 participants that performed the decision-making battery a second time on a follow up 18 months apart on average. Next, we characterised the relationship between the inferred latent cognitive constructs and external measures such as age, IQ, and mental health characteristics. Here, we hypothesized that latent dimensions of decision-making would correlate with self-reported psychological dispositions and mental health symptoms. To test this latter hypothesis, we utilised participants’ derived scores for both general and specific factors of dispositions (Polek et al., 2018) and mental health symptoms (St Clair et al., 2017).

**Table 1:** Cognitive task battery.

| Task (with key reference) | Key constructs assessed | Key individual parameters and descriptives measures. |
| --- | --- | --- |
| A. Go-NoGo task<br>(Guitart-Masip et al., 2012) | Pavlovian biases, i.e. propensity to engage in action in order to obtain rewards and to abstain from action to avoid losses; Motivational power of outcomes; Instrumental learning rate in the appetitive and aversive domains. | 1. Pavlovian Bias;<br>2.-3. Reaction times for action choices in the context of threat vs. opportunity.<br>4. Sensitivity to outcomes.<br>5. General bias for action rather than non-action;<br>6. Motivation-independent, 'irreducible', variability in decision-making; |
|  |  | 7.-8. Learning rates in the appetitive and aversive contexts. |
| B. Approach-Avoidance conflict task<br>(Bach et al., 2014) | Willingness to expose oneself to different levels of risk for the sake of amassing rewards. | 9.-11. Factor-analytic scores summarizing variance over a comprehensive set of behavioural measures in the task.<br>Approximately corresponding to sensitivity to overall level of threat, sensitivity to the time dependency of threat, and overall performance. |
| C. Roulette task<br>(Symmonds et al., 2011)<br>(NB: administered at baseline only) | Baseline taste for gambling<br>Risk-avoidance (preference for outcome distributions of low variance). | 12. Overall preference for gambling over known returns.<br>13. Preference weight for variance, compared to the mean, of an outcome distribution, named 'Economic risk preference';<br>14. Effect of outcome distribution asymmetry (skewness) on preferences.<br>15. Sensitivity to expected value of outcomes. |
| D. Interpersonal-Discounting task<br>(Moutoussis et al., 2016) | Baseline inter-temporal discounting; shift in discounting preferences upon exposure to peers' preferences. | 16. Basic hyperbolic temporal discounting coefficient;<br>17. Relevance of others' observed preferences to the self;<br>18. Discounting taste uncertainty, i.e. uncertainty about one's own tastes in this domain.<br>19. Decision variability over choosing for others<br>20. Irreducible decision noise. |
| E. Information Gathering task<br>(Moutoussis, Bentall, El-Deredy, & Dayan, 2011) | Assessment of whether future decisions will on balance be more advantageous if one gathers more information. | 21. Information Sampling noise, which determines not only decision variability but also effective depth of planning.<br>22. Subjective cost of every piece of information asked for when experimenter imposes no such costs explicitly;<br>23.-24. Ditto if a fixed, external cost-per-step is imposed. |
| F. Multi-round Investor-Trustee task<br>(Fett et al., 2012) | Overall strategies used to elicit cooperation and avoid being exploited by one's anonymous, task partner. | 25. Initial trust, i.e. the amount given by the investor to the Trustee before they have any specific information about them.<br>26. Cooperativeness: Average degree to which Investor and Trustee tended to respond to reductions (or increases) in each other's contributions in kind. |
|  |  | 27. Responsiveness: Average magnitude of responding to the partner's change in contribution. |
| G. Two-Step task (Daw et al., 2011) | Strength of habitual ('model-free') and goal-directed ('model-based') decision-making | 28. Goal-directedness: tendency to shift in decisions as a consequence of a different decision being more advantageous according to the transition probabilities inherent in the task.<br>29. Learning rate<br>30. Perseveration tendency<br>31. Reward sensitivity<br>32. Eligibility trace (propensity of learning to affect not just the current state but also others related to it) |

Crucially, we also characterised the neural circuitry underpinning the latent decision-making factors. To achieve that, we analysed functional connectivity from resting-state fMRI data (rsFC), providing a metric of coupling between blood-oxygen-level-dependent (BOLD) time series from different brain regions or networks (nodes). Patterns of rsFC are known to behave to a large degree as dispositions (Finn et al., 2015) and predict a subject’s cognitive abilities in other domains (Dubois, J., Galdi, P., Lynn, P.K., & Adolphs, R., 2018; Kong et al., 2018; Li et al., 2019; Rosenberg et al., 2015; Smith et al., 2015). We thus asked which connectivity networks predict latent decision-making factors and whether the identified connectivity networks were stable over time.

We found evidence of a single dimension of covariation in the population to which multiple decision-making tasks contributed. This dimension, which we term ‘decision acuity’, reflected speed of learning, ability to heed cognitively distant outcomes, and low decision variability. It showed an acceptable reliability much higher than typical decision-making tasks (Moutoussis et al., 2018) and was associated with distinct patterns of rsFC. Finally, decision acuity was distinct from IQ, as it had a distinct functional connectivity signature and was differentially related to psychological dispositions and symptoms.

## Results

### ‘Decision acuity’ is an important dimension of decision-making

A total of 830 young people aged 14-24 were tested with a battery of tasks assessing fundamental components of decision-making. 349 among them underwent brain functional magnetic resonance imaging at rest, on the same day as cognitive testing, to assess functional connectivity. Scanned participants had no history of neuropsychiatric disorder and were confirmed to be healthy on SCID interview. The Methods section and online Supplement provide further detail.

The decision-making tasks included in the cognitive task battery are described in table 1. Conceptual decision-making constructs overlapped across tasks in the battery, although each task also had a unique focus. Thus we expected participants to show how much they cared about outcomes (reward sensitivity) in almost all tasks. In a similar vein, we expected participants engaging in more sophisticated planning to show increased model-basedness (table 1, task F), better information-gathering (task E), and less temporal discounting (task D). Likewise, we expected participants more capable in interpersonal decision-making to learn more from others (task D) and invest more in benign partners (task F). Finally, we expected those showing excessive risk tolerance to avoid hazards less (task B) and to be less economic risk-averse (esp. in task C). In all, we obtained 32 decision-making measures which we subjected to factor analyses. See Methods for details of the factor-analytic approach, including dimensionality estimation and stability analyses.

Working with the full battery and the larger, baseline sample, we discerned four stable decision-making factors, but only the first loaded on measures from multiple tasks. We named this factor ‘decision acuity’ or *d*, as it loaded negatively on decision variability measures, especially decision temperature, and loaded positively on measures contributing to profitable decision-making, such as low temporal discounting and faster learning rates (Figure 1 and supplemental table S1). Thus, participants with high *d* had low decision variability in economic-risk, information-gathering, Go-NoGo and Two-Step tasks and had fast reaction times and high learning rates in the Go-NoGo task. Note that a decision temperature parameter can always be re-written as the inverse of reward (and/or loss) sensitivity one. Hence the prominent role of negatively-loading temperature parameters in *d* supported our a priori hypothesis that reward sensitivity constitutes an important shared characteristic across tasks. Still in the baseline sample, we confirmed that *d* correlated with profitable decision-making by estimating a measure of aggregate task performance which was based on net points won across tasks, and separate from the components of *d* (Pearson r=0.50, p <1e-10; see Supplement part C for details). Remarkably, *d* predicted this aggregate measure of performance independently from IQ, whereas most of the effect of IQ on performance depended on its shared variance with *d* (performance in tasks and *d* sharing common-method variance being a caveat here).

**Figure 1.**
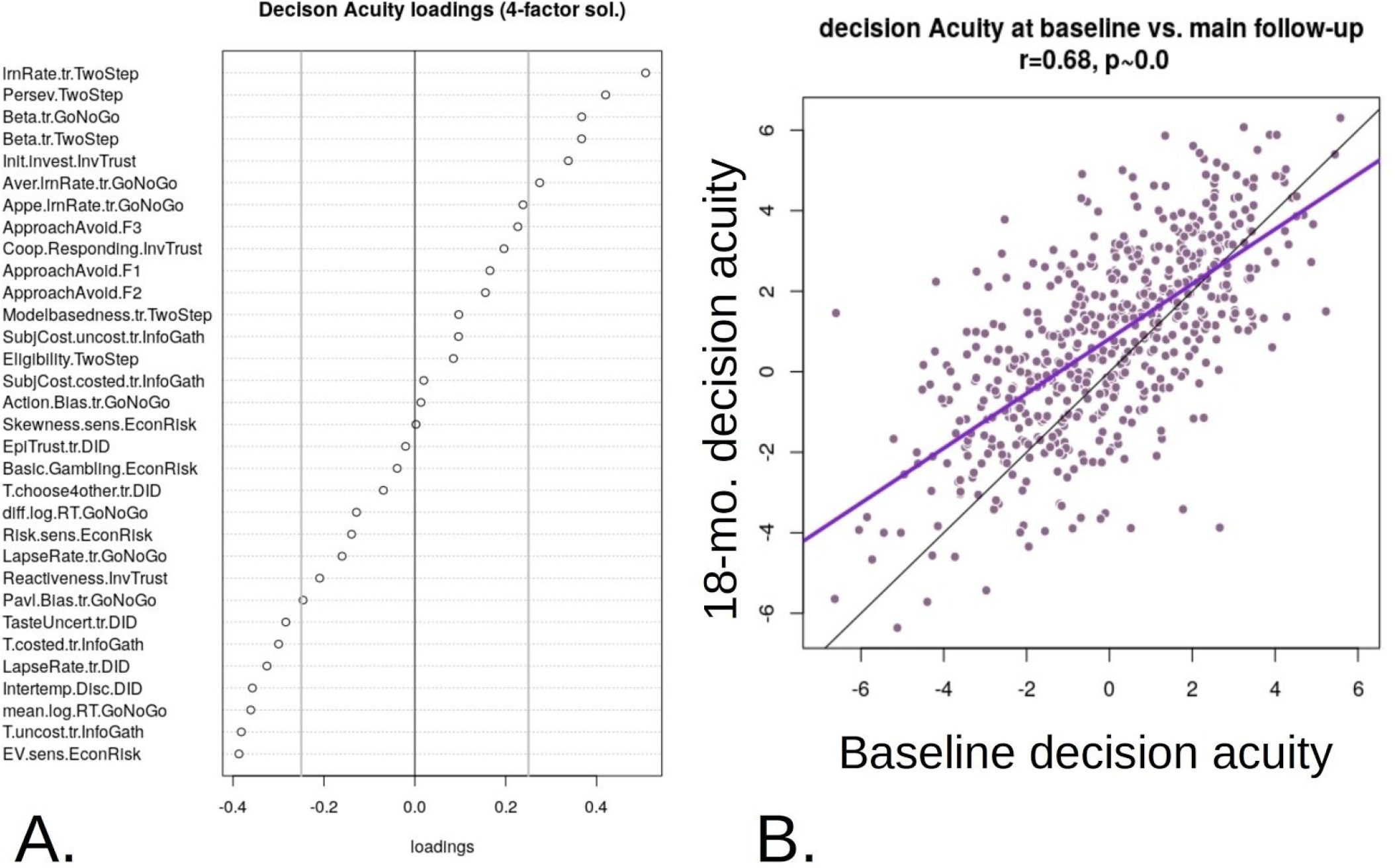
**A.** Decision Acuity common Factor over cognitive parameters, based on the validated 4-factor analysis applied to our whole sample. See supplement table S1 for the key to measure labels. The top half of variables load positively, while grey vertical lines give a visual indication of which measures are important, being the thresholds used for inclusion of variables in the confirmatory analyses **B.** Decision Acuity was strongly correlated between baseline and follow-up, as expected for a dispositional measure. Mauve is regression line, black is identity line.

The other three factors clearly addressed within-task behaviours, rather than hypothesized global decision-making constructs and were thus of peripheral interest here. The second selected the Delegated Inter-temporal Discounting task (D.), the third the Information Gathering task (E.) and the fourth the Economic Risk preference task (C; Figure S2). Over all factors, constituent cognitive measures showed high uniqueness scores, as expected from each task being designed to have a unique focus. 22 of the 32 measures had uniqueness > 80% (Figure 1B).

### Decision Acuity increased with age, follow-up and IQ

We first examined how *d* depended on age, both across and within participants. Linear mixed effects (LME) modelling over baseline and follow-up showed a strong fixed-effect dependence on age (beta=0.24, SE=0.022, p ~ 0.0 (undetectable)). *d* was stable from baseline to follow-up, although slightly less so than WASI IQ (r=0.68, p~0.0 for d; 0.77, p~0.0 for WASI IQ; 95% CI for the difference =−0.135 to −0.044; Fig. 1B) and improved with testing wave (effect size=0.38, p~0.0), but we found no evidence here or in subsequent analyses that its rate of increase depended on baseline age. We then confirmed that both matrix and vocabulary raw IQ (WASI) subscores robustly correlated with *d* (fixed effect betas = 0.088, 0.179, SE= 0.008, 0.018, p ~ 0.0). However inclusion of raw IQ scores did not affect the significance of age as a regressor (age beta=0.121, SE=0.020, p ~ 0.0). Therefore, not only did decision acuity increase with age in our sample, but so did the component that was independent of IQ abilities. IQ subscores and age together accounted for r^2^adj=0.31 of the variance in *d* at baseline.

At baseline, *d* scores for males were higher than females, t-test p=8.6e-5, effect size = 0.27. However, if both IQ subscores and age were entered in LME, the correlation between *d* and self-reported sex was no longer significant. Thus, any uncorrected sex dependence is likely to be due to participant self-selection, that is, amongst males, higher IQ participants volunteered relative to amongst females.

### Mental health factors were specifically associated with Decision Acuity

We next examined the relationship between d and psychological symptoms and dispositions, using scores from published studies of the community sample from whence our participants were sampled (Polek et al., 2018; St Clair et al., 2017). These studies have established that the best descriptions in the symptom and disposition domains were provided by bi-factor models, each comprising a superordinate ‘general factor’ and subordinate ‘specific factors’. Symptoms were described by a general distress factor (a.k.a. ‘p-factor’, (Caspi et al., 2014) and 5 specific factors: Mood, Self-confidence, Worry, Aberrant thinking and Antisocial behaviour. Dispositions were described by a general social functioning factor and 4 specific factors, Social sensitivity, Sensation seeking, Effortful control, and Suspiciousness.

*d* could be significantly predicted by symptoms and dispositions. To test for this, we used LME analysis with participant as random effect, two timepoints of symptoms and decision acuity, and one (baseline) score per participant of dispositions. We first regressed all six symptom scores and five disposition scores against *d*, allowing all to compete to explain variance. We found that amongst symptom scores, *d* was most strongly and negatively associated with the ‘Aberrant thinking’ specific factor, (p=0.0007, bz=−0.19, SE(bz)=0.051). No other symptom factors were significant, (symptom general factor, ‘Distress’, p=0.82, others ranging from p=0.35 to 0.99). *d* significantly related to the general disposition factor, ‘general social functioning’ (p=0.0002, bz=0.36, SE(bz)=0.096). It did not relate to specific dispositions (p ranging from 0.47 to 0.80). We then additionally included raw IQ scores in the LME models. As expected, both raw IQ scores and age significantly predicted *d,* and model fit improved substantially (BIC = 4873 vs. 5083 without IQ). Inclusion of IQ reduced significance of ‘Aberrant thinking’, which draws on schizotypy and obsessionality, to trend level, p=0.074, bz=−0.10, SE(bz)=0.053) but if anything strengthened the significance of ‘Prosociality’ (p=0.0001, bz=0.32, SE(bz)= 0.084). All these analyses also accounted for age as above, and did not benefit from more complex models of age.

### Patterns of brain Connectivity are associated with Decision Acuity differently from IQ

Out of 349 subjects who were scanned at baseline, we discarded baseline scans without acceptable imaging data quality (3), whose ME-ICA denoising did not converge (4), who had a diagnosis of depression (36) or who had excessive motion while scanning (8), leaving 298 baseline scans for analysis. A further three subjects were removed from analyses involving IQ scores as they did not complete the IQ tests, leaving 295 subjects for analysis. A population-average parcellation of brain data was obtained using independent component analysis in our sample, resulting in 168 networks (nodes) *within* each of which activity was highly correlated (see Online Methods for details). Patterns of connectivity *between* nodes were then estimated as partial correlation values, or resting state functional connectivity (rsFC). We then used rsFC values as features in sparse partial least squares (SPLS) analyses, to predict decision acuity and composite IQ. We used cross-validation to prevent overfitting, and predictive accuracy was assessed as Pearson’s correlation coefficient between true scores and values predicted by the model (Figure S4 and Online Methods for details).

Scores for *d* predicted on the basis of functional connectivity, *d*_*pr*′_, significantly correlated with measured *d* controlling for demographic and imaging-related covariates (see methods for details), r=0.145, p<10^−6^. The correlation between measured IQ and IQ predicted on the basis of rsFC using all connections was lower but also significant (r=0.092, p=9e-5).

To interpret the neuroanatomical structure of the predictive model, we first partitioned the nodes into anatomically meaningful ‘modules’ using a community detection algorithm ((Blondel et al., 2008); see Methods), and then asked how well each of these modules predicted *d.* The community detection algorithm clustered the nodes into modules based on the strength of their intrinsic connectivity into disjoint communities to some extent analogous to large-scale functional networks. As shown in figure 2, we obtained the following modules: anterior temporal cortex including the medial temporal lobe (ATC); frontal pole (FPL); frontoparietal control network (FPN); left dorsolateral prefrontal cortex (LDC); medial prefrontal cortex (MPC); orbitofrontal cortex, medial and lateral (OFC); opercular cortex (OPC); posterior cingulate cortex (PCC); posterior temporal cortex (PTC); right dorsolateral prefrontal cortex (RDC); subcortical (SUB); salience network (SAN); somatosensory and motor areas (SMT); visual regions (VIS). We fitted a different SPLS model to the subset of connections involving the nodes in each module, including both intra- and inter-modular connections.

**Figure 2.**
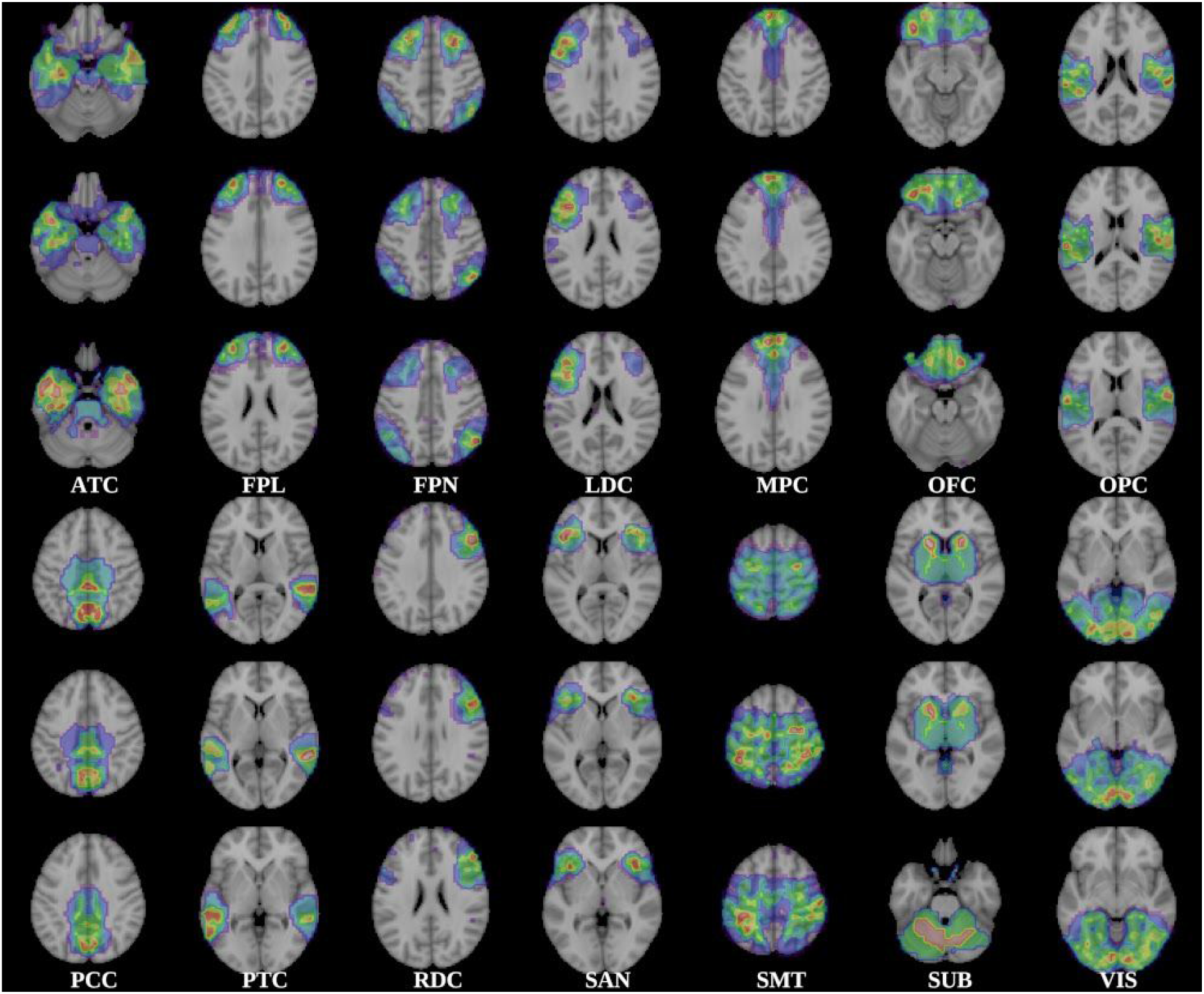
Modules detected by the community structure algorithm. The 168 nodes of the parcellation were clustered in 14 modules with high average rsFC among their nodes. ATC, anterior temporal cortex including the medial temporal lobe; FPL, frontal pole; FPN, frontoparietal control network; LDC, left dorsolateral prefrontal cortex; MPC, medial prefrontal cortex; OFC, orbitofrontal cortex, medial and lateral; OPC, opercular cortex; PCC, posterior cingulate cortex; PTC, posterior temporal cortex; RDC, right dorsolateral prefrontal cortex; SUB, subcortical; SAN, salience network; SMT, somatosensory and motor areas; VIS, visual regions.

The correlation between measured and predicted *d* scores was significant for the FPN, MPC, OFC, OPC, PCC,SMT, and VIS modules after correction for multiple tests (Figure 3A, Table 2), with the strongest correlations for OFC, PCC and SMT. For the PCC and SMT modules, the correlation coefficients exceeded to a small degree the correlation for a model employing all possible connections. This can be explained as a result of feature selection. In the full model it is harder to select just the right features and protect against over-fitting, resulting in a greater penalty in predictive accuracy. On the other hand, the model trained only on a smaller set of features is less likely to overfit. This paradoxical increase in accuracy for a model with less features is known to be stronger when the number of observations is small, relative to the number of features (Chu, Hsu, Chou, Bandettini, & Lin, 2012), which is the case in our dataset. The different modules comprised diverse numbers of nodes but there was no significant association between the number of model features and the correlation between observed and predicted scores (*d*: r=0.356, p=0.193; IQ composite scores: r=-0.158, p=0.574).

**Figure 3.**
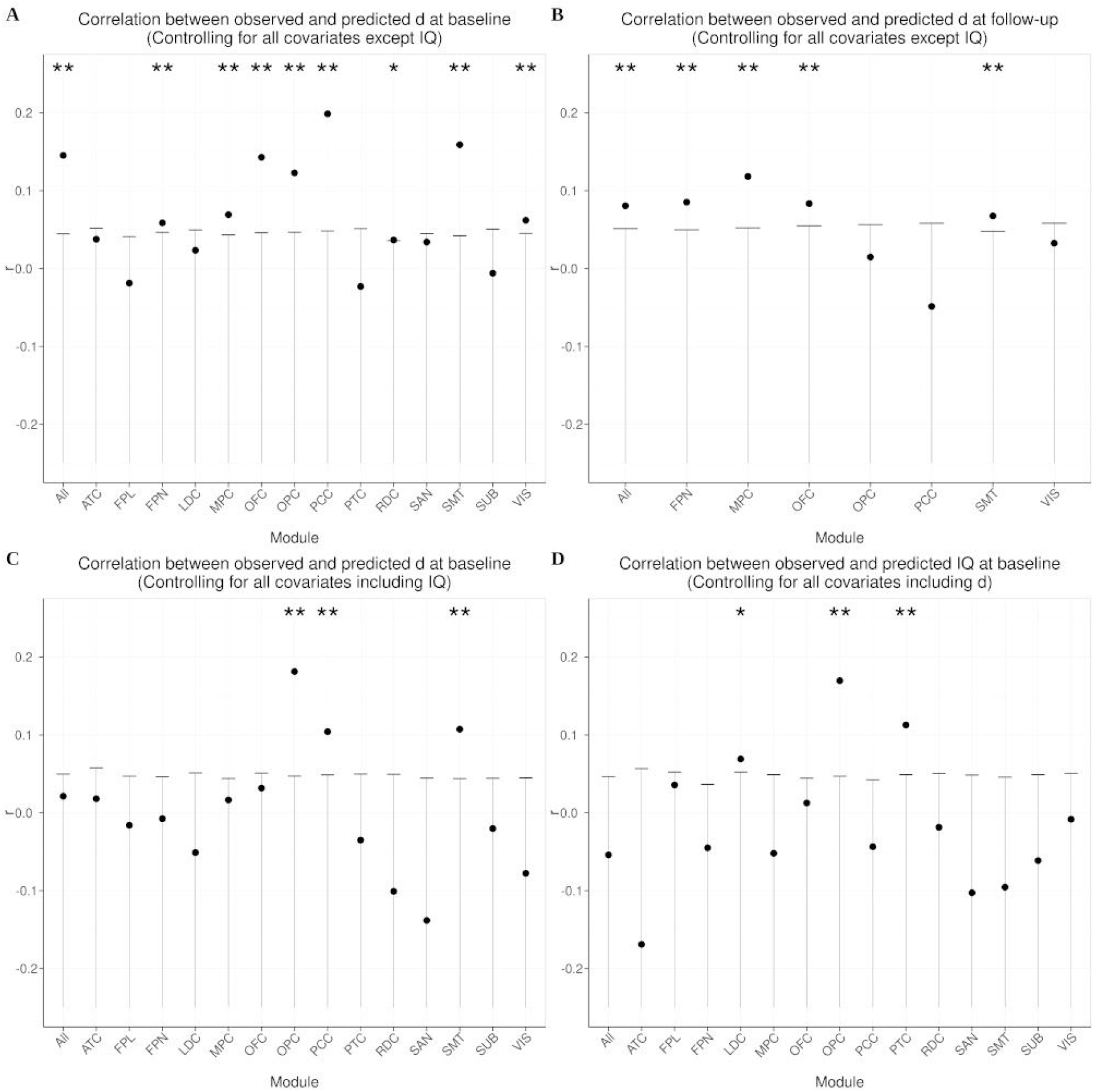
Model predictive performance for each of the functional modules. **A.** Coefficient for the correlation between observed *d* and *d*_*pr*_ predicted by models trained on all connections, and the connections involving nodes in each module. **B.** Correlation between observed d and *d*_*pr*_ predicted by models trained on the baseline data. Only modules for which the prediction was significant at baseline are shown here. All the models included as covariates demographic and imaging-related factors (brain volume, scanning site, head motion; see Online Methods). **C.** As in A., correlation between observed *d* and *d*_*pr*′_, but here additionally correcting for IQ. **D.** Correlation between observed and predicted IQ, but correcting for imaging related factors and decision acuity. In all plots, the leftmost bar corresponds to the model which includes all connections. The whiskers indicate the intervals containing the lower 95 % probability mass for the null distribution, corresponding to one-tailed tests. * significant uncorrected ** significant with FDR correction for the 15 tests. ATC, anterior temporal cortex including the medial temporal lobe; FPL, frontal pole; FPN, frontoparietal control network; LDC, left dorsolateral prefrontal cortex; MPC, medial prefrontal cortex; OFC, orbitofrontal cortex, medial and lateral; OPC, opercular cortex; PCC, posterior cingulate cortex; PTC, posterior temporal cortex; RDC, right dorsolateral prefrontal cortex; SUB, subcortical; SAN, salience network; SMT, somatosensory and motor areas; VIS, visual regions.

**Table 2.**
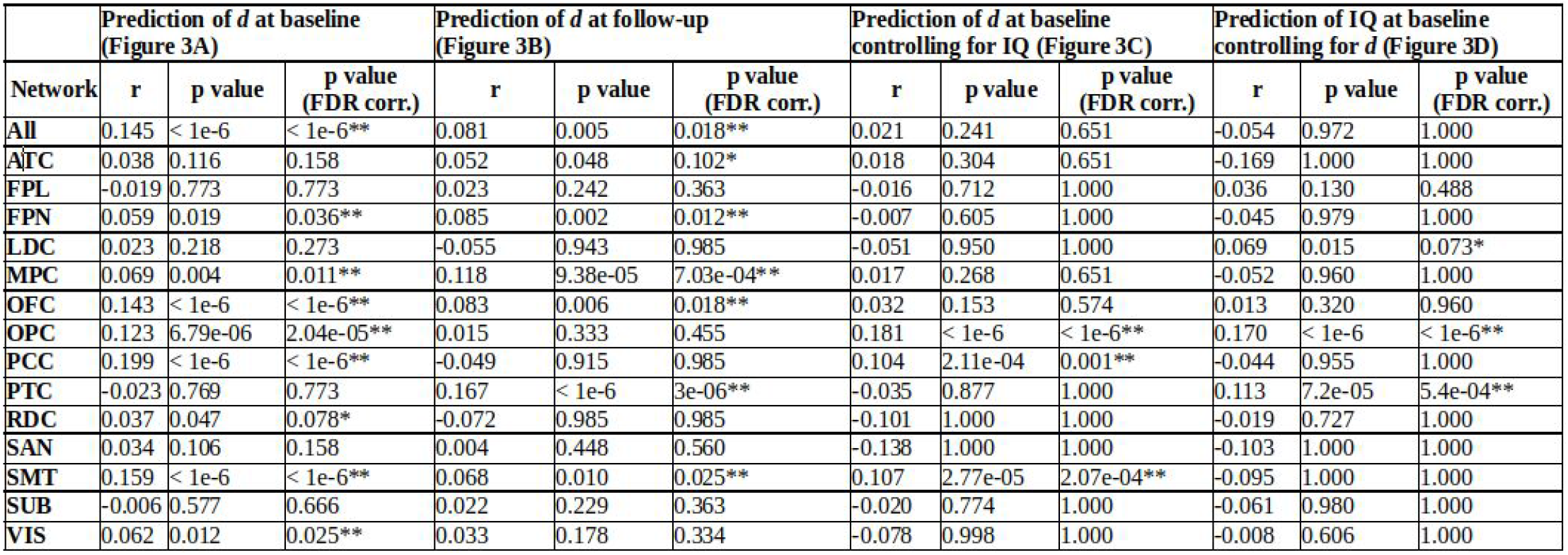
Correlation coefficients between observed and predicted scores, corresponding to the plots in Figure 3. * significant uncorrected ** significant with FDR correction for the 15 tests.

Out of 235 subjects who were scanned at follow-up, adhering to the same criteria as for the baseline data, we discarded those without acceptable imaging data quality (4), whose ME-ICA denoising did not converge (5), and who presented with excessive motion (3), leaving 223 subjects available for analysis. We applied the model trained on the baseline data to the follow-up data (see online methods) for the modules where the prediction was significant at baseline. Importantly, the prediction of a subject at follow-up did not involve their own rsFC baseline data (see online methods), as this would have inflated the estimate of predictive performance. The baseline model predicted significantly the follow-up *d* values based on the follow-up connectivity data when using either all the connections or those with networks in the FPN, MPC, OFC and SMT modules, controlling for demographic and imaging related covariates, and correcting for multiple tests (Figure 3B, Table 2).

In order to assess whether *d* and IQ can be predicted by specific rsFC patterns, or alternatively whether both are underpinned by similar patterns of neural connectivity, we controlled for IQ the partial correlation coefficients between *d*_*pr*_ and *d*, on top of the nuisance covariates previously included. In a complementary manner, we controlled for *d* the partial correlation between *IQ*_*pr*_ and *IQ* (on top of the nuisance covariates). After correction for IQ composite scores, and correcting for multiple comparisons, the correlation between *d* and *d*_*pr*_ remained significant for OPC, PCC and SMT (Figure 3C, Table 2), suggesting that these modules reflect decision acuity over and above their relation to IQ. On the other hand, the correlation between IQ*pr* and IQ was significant for OPC and PTC after controlling for *d* (Figure 3D, Table 2), suggesting that these modules reflect IQ over and above their relation to decision acuity. These analyses demonstrate that decision acuity and IQ have distinguishable and specific signatures in functional connectivity networks: decision acuity taps on the default mode, salience and sensorimotor networks, whereas IQ taps on the salience network but also on temporal networks associated with language processing.

## Discussion

This, to our knowledge, is the first study characterising a distribution of core decision-making measures in an epidemiologically informed sample of adolescents and young adults and relating them to brain function. We found that decision-making performance could be described by a broad construct receiving contributions from multiple domains of cognition. We term this ‘decision acuity’, *d*. In our sample, *d* showed satisfactory longitudinal stability, increased with age and with IQ. *d* also had specific associations with mental health measures, over and above IQ. Importantly, decision acuity showed a temporally stable association with rsFC, involving networks known to be engaged by decision-making processes. Moreover, rsFC patterns associated with *d* and IQ were distinguishable and specific, despite showing some overlap.

Decision acuity had an interpretable structure, conducive to good decision-making. It increased as decision variability lessened, evidenced by its loadings on decision-noise-like parameters across all the tasks that provided such measures. The most prominent such loadings were inverse temperature parameters, also known as reward sensitivities. By definition, high temperature (a.k.a. reduced reward sensitivity) agents care less about relevant outcomes. This supported our hypothesis that reward sensitivity loaded on an important common factor. However, *d* also received substantial contributions from measures that did not directly reflect reward sensitivity, but characterised good decision-making. These included low temporal discounting, fast reaction times, high learning rates, baseline trust in others, low propensity for retaliation, low propensity to show a Pavlovian bias and low lapse rates. Such non-temperature constructs may also be linked to decision variability, albeit less directly.

An interesting interpretation of this pattern is that lower-acuity participants may find it too costly to eliminate computational errors in the fast pace of many tasks. For example, the computations required to make decisions about outcomes far in the future may be hard to perform for low-*d* agents, resulting in discounting-like behaviour. Lapse rates may be understood as ‘floor’ error rates imposed by computational costs. Higher decision variability may also be driven by effective beliefs about the world, for example a belief that over-values exploration. If working out the correct action is too difficult, trial-and-error may be an alternative way to find answers, so this may be a compensatory or adaptation strategy in the face of limited cognitive resources. Overall, the contrast of noise with precision-enhancing measures in this factor is reminiscent of the association between low ability to reach goals and low policy precision in active inference (Friston et al., 2013). The agnostic derivation but interpretable nature of *d* can thus be seen as an example of data-driven ontology (Eisenberg et al., 2019).

High decision acuity was associated with older age, increasing by 0.37 SD over the decade of 14 to 24 years of age, once raw IQ scores were accounted for. This is important as component parameters have been found to have weak or variable relationships with age in this same sample (Moutoussis et al., 2018, 2016). Developmentally, *d* increasing with age may reflect a process whereby adolescents and younger adults get more confident with the outcome of their actions as they age. Next, *d* was associated with psychopathology over and above IQ, specifically increasing with the self-assessed interpersonal competence (‘general social functioning factor’). *d* also decreased with schizotypy/obsessionality traits (‘Aberrant thinking’ factor), but this could be better explained by raw IQ scores. *d* explained a small proportion of the variance in psychopathology, as risk factors often do (Pearson et al., 2015). Overall, though related to IQ, *d* had distinct relationships with mental health measures.

Decision acuity was also associated with specific, distributed patterns of resting-state brain connectivity. The whole brain, connectivity-based predictive model depended on connections spread across the entire brain, implying that *d*, like IQ, depends on more extensive systems than those typically observed for state-tapping tasks in functional imaging studies (e.g. medial prefrontal, dorsolateral prefrontal). Strikingly, the pattern of connections predicting *d* was structured, with connections involving nodes in FPN, MPC, OFC, OPC, PCC, SMT and VIS being most predictive of *d*, irrespective of age and sex. Furthermore, the models trained at baseline on all the features, as well as those restricted on features within FPN, MPC, OFC and SMT, were also predictive of *d* at follow-up, demonstrating the stability over time of the relationship between rsFC in these modules and *d*.

It is unsurprising that decision acuity could be predicted by connections involving MPC and OFC, as these regions are typically recruited by decision-making tasks (Garvert et al., 2015; Padoa-Schioppa & Assad, 2006; Rushworth et al., 2011). Circuits involving these regions receive highly processed sensory information and support goal-directed behaviour by representing subjective value of stimuli and choices. The OFC also supports credit assignment during reward learning (Jocham et al., 2016; Walton et al., 2010) probably by representing the associations between stimuli and outcomes (Boorman et al., 2016; Padoa-Schioppa & Assad, 2006; Stalnaker et al., 2018). Finally, the OFC has also been suggested to support the representation of latent states necessary to navigate decision-making tasks (Schuck et al., 2016; Wilson et al., 2014). Similarly, involvement of the PCC, FPN and SMT is not surprising. Activity in the posterior cingulate cortex has been observed during decision-making tasks and it has been suggested that the PCC monitors the environment to detect transitions to new states (Pearson et al., 2011). Although the frontoparietal circuit has mainly been associated with performance of working-memory tasks (Murray et al., 2017), it has been shown that working memory mechanisms contribute to learning in typical reinforcement learning tasks (Collins et al., 2017; Collins & Frank, 2018). Finally, connections involving motor and somatosensory areas may contribute to adaptive decision-making. For example, in our tasks, motor actions were orthogonalized with respect to choices, and recent work suggests that only the more able decision-makers successfully uncouple motor action and option choice (Shahar, Moran, et al., 2019). Hence, SMT connectivity may be important to achieve this decoupling. Similarly, active suppression of Pavlovian tendencies that can corrupt optimal decision-making may also involve optimal sensorimotor functioning (Cavanagh et al., 2013; Swart et al., 2018).

Our ability to predict decision acuity at baseline when controlling for IQ, as well as IQ when controlling for decision acuity, based on particular connectivity modules demonstrates that both constructs have specific signatures in rsFC. This demonstrates that decision acuity has a neurobiological substrate distinct from that of IQ and further validates the distinctiveness of their association with psychological measures. Although IQ absorbed the predictive ability of the connections within the FPN, the MPC, and OFC, decision acuity tapped on modules within the default mode (PCC), salience (OPC) and sensorimotor (SMT) networks independently of IQ. On the other hand, IQ tapped on the salience network (OPC) too, but also on temporal networks associated with language processing (PTC), consistent with the vocabulary subscale of IQ being heavily reliant on linguistic ability (Axelrod, 2002). Interestingly, connections within the OPC, which encompasses the insula, independently contributed to predicting both decision acuity and IQ at baseline. As part of the salience network, these regions may contribute to modulate the switching between internally and externally directed cognitions (Uddin, 2015).

Decision acuity was related to the mental health indicator, ‘general social functioning’, independently from IQ. This suggests that differences in decision acuity may confer (or indicate) vulnerability to specific psychopathologies. Future studies can usefully build on these observations, as rsFC data can be acquired quickly and does not impose cognitive demands on patients. This endeavour can benefit from advances in computational modelling of cognitive and behavioural data (Huys, Maia, & Frank, 2016), improvements in imaging data collection, processing and modelling (Ciric et al., 2018; Kundu et al., 2017; Todd et al., 2016; Vidaurre, Smith, & Woolrich, 2017), and initiatives to acquire high quality large-scale datasets (Kiddle et al., 2017; Van Essen et al., 2013).

We acknowledge limitations of the present study. We had a retention rate between baseline and follow up of 70%. Although this is acceptable, it meant that our follow-up sample was smaller and we had reduced power to detect longitudinal effects. Although epidemiologically stratified, our sample was a volunteer one, introducing potential self-selection biases. Finally, the reliability and ecological validity of task-based measures would benefit from further improvement.

## Conclusion

We describe a new cognitive construct, decision acuity, that captures global decision-making ability. High decision acuity prominently reflected low decision variability. Decision acuity showed acceptable reliability, increased with age and was associated with mental health symptoms independently of intelligence. Crucially, it was associated with distinctive resting-state networks, in particular in brain regions typically engaged by decision-making tasks. The association between decision acuity and functional connectivity was temporally stable and distinct from that of IQ.

## Methods

### Participants

Participants were invited from a non-clinical community sample until 780 were evenly recruited across 5 age bins (14-16 years,16-18 etc.) and two sexes. Of these, 300 healthy participants were invited for MRI scanning. We supplemented this non-clinical sample with 50 young people recently diagnosed with DSM-5 major depressive disorder. The depressed cohort was excluded from MRI analyses reported here. The study was approved by the Cambridge Research Ethics Committee (12/EE/0250). All participants (and their parents, if less than 16 years old) gave informed consent to participate.

### Decision measures

We used a task battey to assess fundamental aspects of decision-making, namely sensitivity to rewards and losses, attitudes to risk, inter-temporal and reflection impulsivity, pro-sociality and goal-directedness. The battery is presented in table 1 and described in more detail in the supplement. Good performance attracted proportionally greater fees in real money (see Supplement).

Key measures were first extracted from each task according to published methodologies. 830 participants (including all scanned participants) yielded usable data across tasks.

We were interested in whether common factors operated across domains of decision-making. We therefore pre-processed the data to reduce strong correlations among measures within-task, which would otherwise dominate the factor analysis, as is described in the Supplement. In total we formed 32 measures, listed in table 1 and detailed in the Supplement.

### Derivation, validation and psychometric correlates of Decision Acuity

We tailored our analysis to test the hypothesis that around three dimensions of covariation would meaningfully load across decision-making measures, expecting reward sensitivity, risk preferences, goal-directedness and prosociality to be represented in these dimensions. We allowed, however, the data to determine the number of factors in the model. We used an exploratory-confirmatory approach to establish the structure of the factor model using the baseline data. Then, we made use of the longitudinal nature of our sample to test the temporal stability and predictive validity of the key derived measure.

Task measures at baseline only were randomly divided into a ‘discovery’ and ‘testing’ samples. N=416 participants were used for exploratory common factor analysis (ECFA) and 414 were used for out-of-sample testing. We found loadings on the first ECFA factor to vary smoothly across all parameters, and the great majority of loadings to be lower than the conventional threshold of 0.4 (Muthén & Muthén, 2008) (cf. Figure 1). Therefore, for the out-of-sample confirmatory analysis, we allowed for all decision-making items to contribute, recognizing that individual item weights might be poorly estimated, but expecting that the resulting overall scores would be well estimated. We tested this by comparing (i) discovery vs. test samples and (ii) purposeful half-splits of the population with respect to sex and age (see Supplement). Overall, the exploratory analyses suggested that only one common factor, - which we termed ‘decision acuity’, *d* - was relevant to our study questions and that within the range of three to five factors, *d* scores were not sensitive to the exact number of factors. (see supplement). We thus opted for a 4-factor model for all subsequent analyses.

We then tested whether decision acuity as a construct was stable with respect to (i) the random discovery/confirmation split (ii) median-split age and (iii) sex (Supplement B). Finally, we tested for external validity of decision acuity in correlating with (iv) mental health scores for symptomatology and dispositions, using bifactor scores and (v) patterns of functional brain connectivity, as described in Results.

### MRI data acquisition

MRI scans were acquired on three identical 3T whole-body MRI systems (Magnetom TIM Trio; VB17 software version; Siemens Healthcare): two located in Cambridge and one located in London. Reliability of the MRI procedures across sites has been demonstrated elsewhere (Weiskopf et al., 2013). Structural MRI scans were acquired using a multi-echo acquisition protocol with six equidistant echo times between 2.2 and 14.7 ms, and averaged to form a single image of increased signal-to-noise ratio (SNR); TR = 18.70 ms, 1.0 mm isotropic voxel size, field of view (FOV) = 256 × 256, and 176 sagittal slices with parallel imaging using GRAPPA factor 2 in anterior-posterior phase-encoding direction. Resting-state blood-oxygen-level dependent (BOLD) fMRI (rsfMRI) data were acquired using multi-echo acquisition protocol with three echo times (TE = 13, 31, 48 ms), TR of 2420 ms, 263 volumes, 3.8 mm isotropic voxel size, 34 oblique slices with sequential acquisition and a 10% gap, FOV = 240 × 240 mm and matrix size = 64 × 64 × 34. The duration of the functional scan was approximately 11 minutes.

### Connectivity Analysis

The rsfMRI data were denoised with multi-echo independent component analysis (ME-ICA) (Kundu et al., 2017). ME-ICA leverages the echo time dependence of the BOLD signal to separate BOLD-related from artifactual signal sources, like head motion. The functional images were normalized to MNI space by composing a rigid transformation of the average functional image to the participant’s structural image and a non-linear transformation of the structural image to the MNI template, and finally smoothed with a 5 mm full-width-at-half-maximum Gaussian kernel. Following (Smith et al., 2015), group-ICA was applied to the pre-processed fMRI baseline data to decompose it in 200 nodes, 32 of which were identified as artefacts by visual inspection and excluded. The remaining 168 nodes are either confined brain regions or networks formed by regions where BOLD signal time-series are strongly correlated. Multiple spatial regressions against the group-ICA spatial maps were used to estimate time-series for each network and subject, for both baseline and follow-up scans. RsFC matrices (168 × 168 nodes) were then computed using partial correlation with limited L2 regularisation (Smith et al., 2011). All these preprocessing steps were conducted with the ME-ICA toolbox (https://afni.nimh.nih.gov/pub/dist/src/pkundu/README.meica) and the FMRIB Software Library (FSL, https://fsl.fmrib.ox.ac.uk/fsl).

The obtained rsFC values were used as features in a sparse partial least squares (SPLS) model to predict two outcome measures of interest (decision acuity and IQ composite scores). SPLS ((Chun & Keleş, 2010); ‘spls’ R library, https://cran.r-project.org/web/packages/spls/) is a multivariate regression model that simultaneously achieves data reduction and feature selection. It has application in datasets with highly correlated features and sample size much smaller than the total number of features, as was the case in the present study. SPLS models are governed by two parameters (number of latent components and a threshold controlling model sparsity) that were adjusted using a nested cross-validation scheme (i.e. using data in the training dataset only) with 10 folds (supplement Figure S4).

Predicted scores were estimated by 20-fold cross-validation repeated 5 times. For each training-testing partition we performed the following steps. To elucidate whether the predictions were driven by rsFC values independently of age, sex or covariates of no interest (see below), we fitted a linear model to the training dataset and regressed out from the target variable (in both training and testing datasets) age, sex and their interaction as well as brain volume, scanning site and head-motion-related parameters. Head motion is known to originate spurious correlations that bias connectivity estimates and therefore (besides the ME-ICA preprocessing explained above) we regressed out average framewise displacement (FD), a summary index of the amount of in-scanner motion (Power, Barnes, Snyder, Schlaggar, & Petersen, 2012), and the degrees of freedom resulting from the ME-ICA denoising, which may differ across subjects depending on how much nuisance variance is removed from their data. As an additional control for head motion, subjects whose mean FD was above 0.3 mm were not included in the analysis. We also standardized both training and testing data with respect to the mean and standard deviation of the training data (separately for each feature). As a first step to filter out uninformative features and speed up computations, only those significantly (p < 0.05) correlated with the outcome variable in the training dataset were entered in the SPLS model. We then used a bagging strategy where data were resampled with replacement 200 times and as many SPLS models were fitted to the resampled datasets, and their feature weights averaged to produce a final model. The purpose of this step was 1) to improve the generalizability of the final average model and 2) to allow estimation of the stability of the feature weights selected. The final, average model was used to compute the predicted scores for the testing partition. The same procedure was repeated for all folds to obtain one predicted score for each subject, where the predicted score for each participant depended only on data from other subjects in the sample. These procedures were implemented with R (https://www.r-project.org/) and MATLAB (https://www.mathworks.com).

### Network node community structure

To enhance our understanding of the anatomical distribution of the predictive connections, we performed a ‘virtual lesion’ analysis (Dubois, J. et al., 2018), which entails assessing the performance of the model when it is trained only on subsets of connections instead of the full ensemble. First, we partitioned the set of nodes into disjoint modules or communities (to some extent analogous to large-scale functional networks (Smith et al., 2009)) formed by nodes which displayed high connectivity among them but lower connectivity with nodes in other modules. We obtained the community structure directly from our dataset instead of relying on previous partitions that have been derived from adult connectomes (Ito et al., 2017; Power et al., 2011) (Ito et al., 2017; Power et al., 2011), because brain connectivity of adolescents and adults is known to differ (Fair et al., 2009).

To produce the partition, we averaged the baseline rsFC matrices across participants and removed negative entries. The resulting matrix was submitted to the Louvain community detection algorithm for weighted graphs (Blondel, Guillaume, Lambiotte, & Lefebvre, 2008) and this partition was refined using a modularity fine-tuning algorithm (Sun, Danila, Josić, & Bassler, 2009). Since the algorithm is not deterministic, it was applied 100 times and the results gathered in a nodes x nodes consensus matrix that indicates the frequency by which the corresponding node pair was assigned to the same module. The consensus matrix was partitioned repeatedly until convergence. The algorithm depends on a parameter γ that controls the resolution (which determines the ensuing number of modules). We adjusted this parameter to maximize the normalized mutual information between solutions at different resolutions. The optimal value of γ ensures the most stable partitioning and in our dataset (γ=2.7) led to a solution with 14 modules, a number that yielded interpretable modules and is on par with the cardinality used in previous studies. These analyses are similar to those reported in (Geerligs, Rubinov, Cam-CAN, & Henson, 2015) and were performed with the Brain Connectivity Toolbox ((Rubinov & Sporns, 2010), www.brain-connectivity-toolbox.net) for MATLAB. Having parcellated the connectome in the 14 modules, we trained the prediction model for each one of them using only connections implicating nodes in that module (i. e. either connections among nodes in the module or connections between nodes in the module and the rest of the brain). We employed the same module decomposition in the analysis concerning the follow-up dataset.

### Predictive performance

We assessed predictive performance as the Pearson correlation coefficient *r* between measured *d* and (cross-validated) predicted *d* (*d*_*pr*_), averaged across repetitions of the cross-validation splits. After Fisher transformation, the null distribution of *r* should follow a zero-centered Gaussian distribution. In order to appraise significance, we estimated the variance of this distribution by generating 30 random permutations of the target variable (Winkler et al., 2016) and repeating the model-fitting procedures mentioned above, separately for each fold. We then derived p-values for the observed *r* from the estimated null distribution. We assessed predictive performance for a model based on the full set of connections, as well as for models trained on the subsets of connections corresponding to the modules described in the previous subsection. To demonstrate that the relationships between connectivity and decision acuity were stable over time and replicate, we used the model estimated at baseline to predict *d* based on the follow-up rsFC data for modules that were significant at baseline. Given that the data at baseline and follow-up are not independent, we kept the same cross-validation fold structure in both datasets, so that the prediction of a subject at follow-up did not involve their own rsFC baseline data, as this would have inflated the estimates of predictive performance at follow-up.

### Connectivity patterns predictive of *d* vs IQ

For imaging analyses, we derived a composite score of IQ by averaging standardized vocabulary and matrix IQ subscores, rather than using the standardized WASI score, because of two reasons. First, we wanted analyses involving both age and IQ to have a straightforward interpretation where IQ represents a measure of raw ability, as opposed to age-standardized ability, and explicitly test for age-dependence separately. Second, we found evidence (Results) that our sample was different from the original on which standardised scores were derived, and hence the standardisation procedure might be invalid. Next, we trained models both on the complete set of connections and the subsets corresponding to the individual modules to predict the IQ composite scores, as we had done previously to predict *d*, yielding IQ_*pr*′_, and assessed predictive performance for each of the modules separately. To compare the connectivity patterns that were predictive of *d* with those predictive of IQ, for each of the modules we assessed the partial correlation between *d* and *d*_*pr*_ when controlling for *IQ*, and the partial correlation between *IQ* and *IQ*_*pr*_ when controlling for *d*. In all these analyses we corrected for age, sex and imaging-related confounds as above.

## Supporting information

On-line Supplement

## Acknowledgements

The Wellcome Trust funded the “Neuroscience in Psychiatry Project” (NSPN). All NSPN members (Supplement section E) were supported by the Wellcome Strategic Award, ref 095844/7/11/Z. Ray Dolan is supported by a Wellcome Investigator Award, ref 098362/Z/12/Z. Statistical analyses were enabled by resources provided by the Swedish National Infrastructure for Computing (SNIC) at the HPC2N center, partially funded by the Swedish Research Council through grant agreement no. 2016-07213. The Max Planck – UCL Centre for Computational Psychiatry and Ageing is a joint initiative of the Max Planck Society and UCL. DRB is supported by funding from the European Research Council (ERC) under the European Union’s Horizon 2020 research and innovation programme (Grant agreement No. ERC-2018 CoG-816564 ActionContraThreat). MM and DRB receive support from the National Institute for Health Research (NIHR) UCLH Biomedical Research Centre.

