## Supplementary material for "A generic decision-making ability predicts psychopathology in adolescents and young adults and is reflected in distinct brain connectivity patterns": On-line Supplement

#### Supplementary Methods

##### Participants

Participants were sampled from a pool of c. 2400 community-dwelling young people and formed a 'cognitive cohort', as mentioned in the main Methods. Participants were contacted at random from 5 age bins (14-16,16-18 etc.), until each recruited age bin had approximately equal proportions of females and males. The proportion of non-white-English youngsters in our study was within 10% of that of the most recent census. Significant neuropsychiatric problems were screened out by self-report, and recruitment sources were selected for the sample to be as representative as possible of the healthy population (Kiddle et al., 2017). We continued to invite people from the larger pool into the cognitive cohort, until our target number of 780 'cognitive' participants was completed. Of these, 300 were invited for MRI brain scanning. They were equally distributed in the 5 age bins and two sexes as above. In addition, they were screened for absence of a history or presence of mental health disorder, neurological or major health problem, or learning disability. Initial screening was by self-report but was confirmed by SCID-II interview and IQ testing. We supplemented this non-healthcare-seeking sample with 50 young people recently diagnosed with DSM-5 major depressive disorder. Thus, the main sample was representative of the healthy wider population, but a smaller depression group was also analysed to test whether the structure of decision-making and the relevant brain measures identified in the healthy population also extended to this health-seeking group. The depressed cohort was excluded from MRI analyses reported here.

Participants (and their parents, if less than 16 years old) gave informed consent to participate in the study. The study was approved by the Cambridge Ethics Committee (12/EE/0250).

##### Cognitive Task Battery

This assessed fundamental aspects of decision-making, namely sensitivity to rewards and losses, attitudes to risk, inter-temporal and reflection impulsivity, pro-sociality and goal-directedness. The battery is briefly presented in table 1 and described in more detail in the supplement (A, 'Decision-making tasks'). Each task, with the exception of an Information Gathering task (IGT), was an abbreviated version of a task described in published work and summarized in table 1. The IGT was a serial information sampling task adapted more extensively than other tasks for the purposes of this study from (Lincoln, Ziegler, Mehl, & Rief, 2010; Moutoussis et al., 2011).

Tasks lasted 8-30 minutes each, giving an overall duration of 2 ½ – 2 ¾ hours, including one obligatory break and as many extra between-task breaks as the participant asked for. Good performance attracted proportionally greater fees in real money (see Supplement A).

#### Decision measures

Key measures were extracted from each task according to published methodologies. These appear in table 1 and further details are found in Supplement A. 830 participants (including all scanned participants) yielded useable data across tasks.

We were interested in whether common factors operated across domains of decision-making, which meant making a judgement about the variables to include in the factor analysis. Inspection of the bivariate correlation matrix revealed a small number of high inter-correlations, especially within specific pairs of measures. For example, reaction times for the two different types of 'action trials' in the go-nogo task, i.e. 'go-to-avoid-loss' and 'go-to-win', were highly correlated, whereas it was the difference between reaction times in these trial types that was of great interest for decision-making. We therefore re-expressed each of these pairs in terms of their mean and difference, which eliminated these correlations. The Approach-Avoidance task potentially provided many measures with complex inter-correlations. Here we used a separate, within-task factor analysis to reduce correlations (see Supplement A) and used the factor scores of a validated 3-factor solution as input variables in the current analysis. In total we selected 32 measures, listed in table 1.

##### Derivation, validation and psychometric correlates of Decision Acuity

We tailored analysis to test the hypothesis that a few (around three) dimensions of covariation would meaningfully load across decision-making measures, expecting reward sensitivity, risk preferences, goal-directedness and prosociality to be represented in these dimensions. We allowed, however, the data to determine the number of factors in the model. We used an exploratory-confirmatory approach to establish the structure of the factor model using the baseline data. Then, we made use of the longitudinal nature of our sample to test the temporal stability and predictive validity of the key derived measure.

Task measures at baseline only were first transformed to near-normal marginal distributions using logarithmic or power-law transforms, imputed for the small percentage of missing values using the R package 'missMDA', then randomly divided into a 'discovery' and 'testing' samples. N=416 participants were used for exploratory common factor analysis (ECFA) and 414 were used for out-of-sample testing. We found loadings on the first ECFA factor, likely to be most important, to vary smoothly across all parameters, and the great majority of loadings to be lower than the conventional threshold of 0.4 used to construct structural equation models for confirmatory FA (Muthén & Muthén, 2008). Items had high uniqueness, as expected. These results were much like the final total-sample FA illustrated in Figure 1. Therefore, rather than claim that certain decision parameters were important and others were not in providing a measure of the underlying latent variable, we allowed for all decision-making items to contribute, recognizing that individual item weights would be poorly estimated, but expecting that the resulting overall scores would be well estimated. We tested this by comparing (i) discovery vs. test samples and (ii) purposeful half-splits of the population with respect to sex and age (see Supplement B). The exploratory analysis furthermore suggested that our objective need not be to determine a ground-truth number of factors, as higher order factors were dominated by single tasks and hence were of no interest here. Our criterion for including higher-order factors then was whether higher dimensional models were likely to result in better score estimates for the low-order factors, which were of interest. Using three to five factors fulfilled these criteria, and indicated that only the first factor - which we termed 'decision acuity', *d* - was relevant to our study questions. Within the range of three to five factors, *d* scores were not sensitive to the exact number of factors, either in the discovery sample or from discovery to testing samples (see supplement). We thus opted for a 4-factor model for all subsequent analyses.

We then tested whether decision acuity as a construct was stable with respect to (i) the random discovery/confirmation split (ii) median-split age and (iii) sex using the baseline dataWe examined how closely scores for a certain subgroup (say, younger people) based on ECFA of the group itself agreed with scores for the same individuals based on FA weights derived from the opposite (e.g. older) group, described in Supplement B. Finally, we tested for external validity of decision acuity in correlating with (iv) mental health scores for symptomatology and dispositions, using bifactor scores and (v) patterns of functional brain connectivity, as described in Results.

The follow-up battery did not contain one of the baseline tasks, and had minor differences (but the same derived parameters) for two further tasks. In order to perform longitudinal analyses, we adopted a conservative approach, estimating a measure of decision acuity based on the final stage of the baseline analysis, but retaining only the weights for the six tasks that were assessed longitudinally. We checked that this more approximate measure adequately captured individual variability of the baseline sample, which was the case (r=0.98, p undetectable) and therefore used in the longitudinal analysis baseline scores derived from these six tasks. We then derived the follow-up decision acuity estimates as follows. We first applied the same approximate-gaussianization transforms to each follow-up measure. Next, we z-scored each follow-up measure using the mean and standard deviation of the respective (transformed) baseline measure. Finally, we applied the weights for these 6 tasks derived from the baseline factor analysis. Thus, we follow-up measures of decision acuity had exactly the same structure as baseline and could be used to compare absolute changes in this measure.

For the longitudinal analysis, we used a linear mixed effects approach. Developmental time in this accelerated longitudinal design is represented both by age-at-recruitment, and by the time interval between test waves. Both recruitment procedures and development itself may mean that these two measures of developmental age may in practice affect our dependent variables differently. We therefore first checked if LME modelling over baseline and follow-up with age as a random effect, in addition to a random intercept for each participant, improved model fit. In fact, it worsened model fit (BIC = 5974.5; logLik = -2965.512; vs. BIC=5960.0, logLik= -2965.529), so we did not include age as random effect in further analyses. In further analyses involving IQ, we used the raw matrix and vocabulary WASI IQ subscores and modelled age explicitly, rather than use standardized IQ subscores. This is because we noticed that the standardized WASI total IQ in our sample was associated with age (r Pearson=0.135, p=0.00011, $r^2$=0.017) at baseline. This indicates that our sample had a different age dependence of IQ scores than the reference one (Axelrod, 2002). Therefore, we regressed *d* for raw IQ subscores while covarying for age, in effect accounting for variation in IQ ability independent of whether this was due to age or self-selection.

#### A. Decision-making tasks

We selected seven tasks tapping fundamental decision-making with evidence linking them to both mental health symptoms and neural mechanisms (table 1 in main text). First, a Go-NoGo task (Guitart-Masip et al., 2011) provided measures relevant to sensitivity to rewards and Pavlovian bias. Second, an approach-avoidance task measured the balance of seeking rewards vs. avoiding losses (Bach et al., 2014). This is likely to be relevant to everyday risk-taking by young people. Third, a risk preference task (Symmonds, Wright, Bach, & Dolan, 2011) complemented this, focusing on widely accepted economic measures of risk-taking (Bach et al., 2020; Rigoli et al., 2016). Fourth, we assessed inter-temporal discounting, learning about the preferences of others and finally peer influence (Moutoussis, Dolan, & Dayan, 2016; Nicolle et al., 2012). Discounting has been shown to be important in a range of psychiatric disorders (Bickel, Jarmolowicz, Mueller, Koffarnus, & Gatchalian, 2012) and so are issues of thinking about others (Sripada et al., 2009) and peer influence (Kerr, Van Zalk, & Stattin, 2012). Fifth, we included an information gathering task (Moutoussis, Bentall, El-Deredy, & Dayan, 2011)  as this has been consistently shown to be relevant to psychotic symptoms (Lincoln, Ziegler, Mehl, & Rief, 2010)  as well as the fundamentals of decision-making (Dayan, 2014). Sixth, a Trust Task was used as a measure of complex social cognition especially relevant to disorders of interpersonal function (Fett et al., 2012; King-Casas et al., 2008). Seventh, one of our most important tasks assessed the role of habitual vs. planful mechanisms in decision-making (Daw, Gershman, Seymour, Dayan, & Dolan, 2011). The battery was implemented using matlab (MATLAB, 2012) using the Cogent toolbox (see Acknowledgements). Trained research assistants directed the participants through the battery.

In terms of remuneration, participants received a flat fee but were also (truthfully) told that they would be paid extra according to their earnings in the tasks. They were informed that there would be a substantial amount of luck in each task, but those who completed the tasks carefully would expect to earn about 2.5 pounds extra per task. Participants did not see earnings for each trial, because tasks differed greatly in their delivery and we did not want to display varying amounts of money to avoid additional Pavlovian motivational effects. Instead, participants were told that 'roughly, each good decision in each task is worth approximately the same', a statement which provided a reasonable reflection of the true state of affairs. The sole element of deception in the battery was that during the interpersonal tasks participants were told that their play partner was a peer, whereas in reality it was a computer agent. However these agents were simulating as closely as possible the performance of healthy people who had the same demographics as the participants. Participants were debriefed at the end of all testing.

Earnings were added to their compensation for the day's testing, except for the Interpersonal-Discounting task. Here, participants were paid at one of their chosen delays, randomly chosen from all the trials in the task,  if they chose a larger but delayed payment. This was paid in Amazon vouchers.

The order of the tasks was subject to constrained randomization. We first piloted the battery in 15 participants, of whom we asked detailed feedback as to how interesting and how tiring they found each task, as well as free-form comments. On the basis of this we avoided putting the more tiring or less interesting tasks near the end of the battery, in order to minimize the effect of fatigue. This resulted in eight different task sequences, one of which was given at random to participants. After the first 40 participants were recruited we performed an interim analysis to compare performance in this battery of shortened tasks as compared to the full-length versions. Performance in each task showed followed the pattern of performance in the original, except the Two-Step task. Here participants as a group showed only just-detectable goal-directed decision-making. As this would greatly reduce the task's usefulness we improved the pre-task training and instructions and discarded this first ~10% of data for this task, with satisfactory results.

Importantly, the sample of 830 participants which performed the cognitive tasks included 780 who were healthy at the point of testing, but also 50 of their peers diagnosed with DSM-5 major depressive disorder. As mental health symptoms were under-represented in the healthy cohort by construction, we decided a priori to enrich the cohort with individuals with depression to increase sensitivity to capture relationships with symptoms. In the event, however, decision acuity was not related to general-distress or mood symptomatology. Post-hoc, we checked whether excluding the depression participants affected the factor analysis, and consistent with the insensitivity of decision-acuity to mood, we found that it made no difference of note. We therefore report the factor-analyses of the entire sample, as planned a priori.

**Human Approach-Avoidance task**

Here we describe how the task described in Bach and coworkers (2014) was adapted for the purposes of this study. Because of time constraints we reduced the number of threat contexts from three to two, which we call two 'predators' corresponding to low and high threat. Also, different from the previous study, epoch duration did not depend on threat level. That is, an epoch ended after a random duration, independent of whether the predator woke up or not. Finally, the number of epochs was reduced to 1/3 of the original, so that the task took about 23 minutes to complete.

Based on the previous work (Bach et al., 2014) , we collected a large number of behavioural descriptive measures and performed an exploratory factor analysis of these (substantially correlated) measures. We found that the first three factors could be meaningfully interpreted in decision-making terms, namely as sensitivity to the level of threat in the environment ('threat sensitivity'), sensitivity to features increasing probability of loss within an environment ('loss sensitivity') and measures of overall performance ('performance'). As might be expected, this third 'performance' factor loaded more highly in $d$ (Figure 1) but still did not exceed the threshold of 0.25 that we used for inclusion in confirmatory analyses (below).

**Information-Gathering task**

The 'cover story' and graphics of the Information Gathering task were adapted from the work of Lincoln and coworkers (Lincoln, Peter, Schafer, & Moritz, 2010; Lincoln, Ziegler, et al., 2010). On the basis of previous work (Moutoussis et al., 2011) we reduced the maximum number of samples of information per trial in order to increase the impact of the approaching end (urgency). We first presented participants with an uncosted-information gathering version of the task for 10 trials. This had deliberately non-specific instructions to maximise the chance that participants would bring their own, subjective cost structure to bear and because, somewhat unexpectedly, such uncosted, scarce-instruction versions of the task has produced some of the most consistent results in clinical and subclinical samples. We then presented them with 10 trials with more specific instructions. Participants started with 100 points and had to pay 10 points for each item of information they requested. We employed a maximum-likelihood fit of the bayesian-observer model from (Moutoussis et al., 2011).

**Investor - Trustee task**

In order to analyse the Investor-Trustee task we adapted the measures described by (Fett et al., 2012). Following these researchers, we considered whether participants increased or decreased their offer at each move in response to observing their partner increase or decrease theirs. However we considered the fractional change in contribution, i.e. the change in the fraction of play-money that could have been given. This entails the hypothesis that each player considers the other as 'messaging' them from a baseline of their financial means, not in absolute terms. We then considered the vector in the 2-dimensional space of (fractional-change-of-Investor by fractional-change-of-Trustee) formed for each round of play. We classified this in the same way as Frett et al. (retaliating, repairing, honouring, disrupting) as the angle between the vector and the change-of-Investor axis increased from -180 to 180 degrees. Again using the (rather crude) approximation that strategy remains the same throughout the 10 rounds of the game, we added the vectors for each of the rounds to determine the character of the game as a whole. The orientation of the resultant vector characterises the whole exchange – both Investor (our participant) and Trustee (the computer). As all investors played the same computer program, this vector can be seen as the type of exchange that the participant elicited.

In the event, orientations showed a clear bimodal distribution, either around zero degrees (an exchange based on coaxing the Trustee) or around $-3\pi/4$ . The latter represents an exchange where each party is responding to the other's reduction in contribution with their own reduction. We might speculate that participants attempt to signal 'if you won't be generous, I won't either'. The two-cluster distribution could in turn be fitted reasonably well with a single straight line spanning retaliatory to coaxing exchanges. The 'trust building' index in tables 1 and 2 corresponds to the participant's position along this line.

| | |
|---|---|
| lrnRate.tr.TwoStep | **Learning rate, Two -step task (transformed)** |
| Persev.TwoStep | **Perseveration parameter, Two-step task** |
| Beta.tr.GoNoGo | **Inverse temperature, Go-NoGo task (transformed)** |
| Beta.tr.TwoStep | **Inverse temperature, Two -step task (transformed)** |
| Init.Invest.InvTrust | **Initial investment in partner, Investor-Trustee task** |
| Aver.lrnRate.tr.GoNoGo | **Aversive learning rate, Go-NoGo task** |
| Appe.lrnRate.tr.GoNoGo | Appetitive learning rate, Go-NoGo task |
| ApproachAvoid.F3 | 'Performance factor', Approach-Avoidance task |
| Coop.Responding.InvTrust | Degree of cooperative responding, Investor-Trustee task |
| ApproachAvoid.F1 | 'Sensitivity to overall threat level', Approach-Avoidance task |
| ApproachAvoid.F2 | 'Sensitivity to increasing hazard', Approach-Avoidance task |
| Modelbasedness.tr.TwoStep | Model-basedness, Two-step task (transformed) |
| SubjCost.uncost.tr.InfoGath | Subjective cost of samples, uncosted Info. Gathering (transf.) |
| Eligibility.TwoStep | Eligibility trace parameter, Two -step task |
| SubjCost.costed.tr.InfoGath | Subjective cost of samples, costed Info. Gathering (transf.) |
| Action.Bias.tr.GoNoGo | Bias towards action, Go-NoGo task (transformed) |
| Skewness.sens.EconRisk | Sensitivity to outcome skewness, Econ. preference task |
| EpiTrust.tr.DID | Epistemic trust parameter, delegated discounting (transf.) |
| Basic.Gambling.EconRisk | Overall preference for gambling, Econ. preference task |
| T.choose4other.tr.DID | Variability of choices-for-other, delegated discounting (transf.) |
| diff.log.RT.GoNoGo | React. time diff. between conditions, Go-NoGo task (transf.) |
| Risk.sens.EconRisk | Risk aversion, Econ. preference task |
| LapseRate.tr.GoNoGo | Lapse rate, Go-NoGo task (transformed) |
| Reactiveness.InvTrust | Reactiveness to other's offers, Investor-Trustee task |
| Pavl.Bias.tr.GoNoGo | Pavlovian bias, Go-NoGo task (transformed) |
| TasteUncert.tr.DID | **Taste uncertainty, delegated discounting (transf.)** |
| T.costed.tr.InfoGath | **Decision temperature, costed Info. Gathering (transf.)** |
| LapseRate.tr.DID | **Lapse rate, delegated discounting (transformed)** |
| Intertemp.Disc.DID | **Temporal discounting, delegated discounting (transf.)** |
| mean.log.RT.GoNoGo | **mean log-Reaction Time, Go-NoGo task** |
| T.uncost.tr.InfoGath | **Decision temperature, uncosted Info. Gathering (transf.)** |
| EV.sens.EconRisk | **Sensitivity to expected value of outcome, Econ. preference task** |

**Table S1** Key to the labels of cognitive measures, in the order of loading onto decision acuity. Green - load positively; Blue - load negatively; Bold - exceed 0.25 in loading.

### Supplementary Analyses

#### B. Factor analysis and validation of Decision Acuity

##### B1. Exploratory - Confirmatory analyses

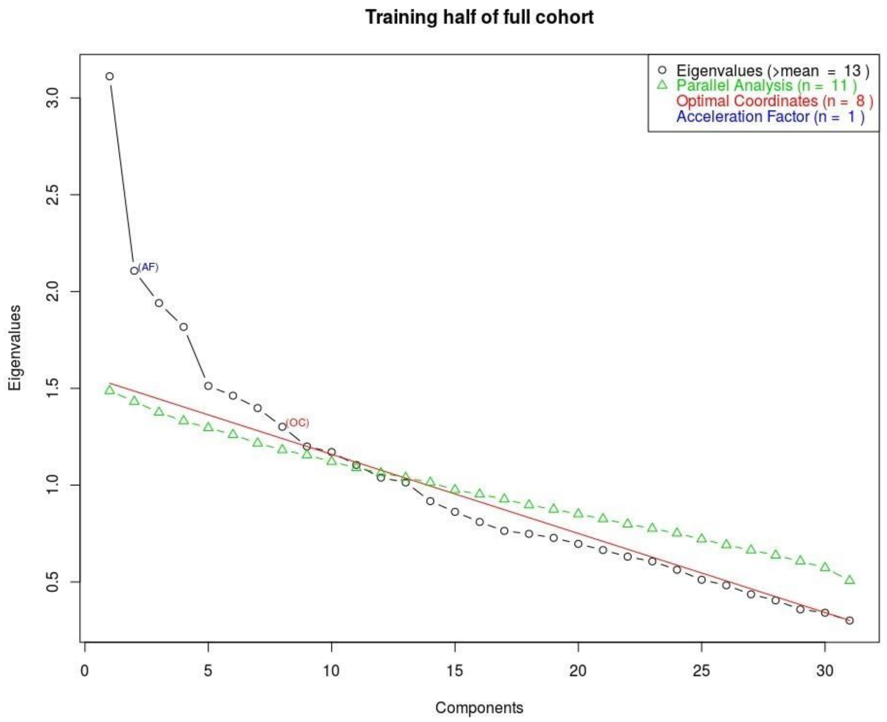


**Figure S1** Parallel analysis to determine optimal number of factor-analytic components for the 'Discovery dataset', N = 416, and 32 variables.

The maximum number of factors for the exploratory-confirmatory analysis was 8, estimated by parallel analysis (Figure S1). The 8 factors estimated from the ECFA were then tested on the confirmation dataset using structural equation modelling in R (Fox, 2006). Criteria of under-determination, Bayesian Information Criterion (BIC), and comparative fit index (CFI) were used to compare including an increasing number of factors. In the confirmatory factor analysis, a threshold of 0.25 was adopted as very few loadings on the first factor exceeded the conventional threshold of 0.4 (See Figure 1 in the main text). Considering the test set of 414 participants only, we found that model fit as indexed by the BIC and CFI improved from 1 to 4 factors. However, a 5 factor and more complex model fits did not converge on the test set. According to these criteria we considered a model of four factors to be most parsimonious and robust. However, as the threshold of 0.25 is not uniquely defined, and as splitting the data set into discovery and testing means that a limited number of data was available to test high-dimensional models in a confirmatory mode, we informed our choice of factor-analytic dimensionality by the stability of the latent variable scores to which different FA models gave rise to. This is described in the 'Stability analysis' below.

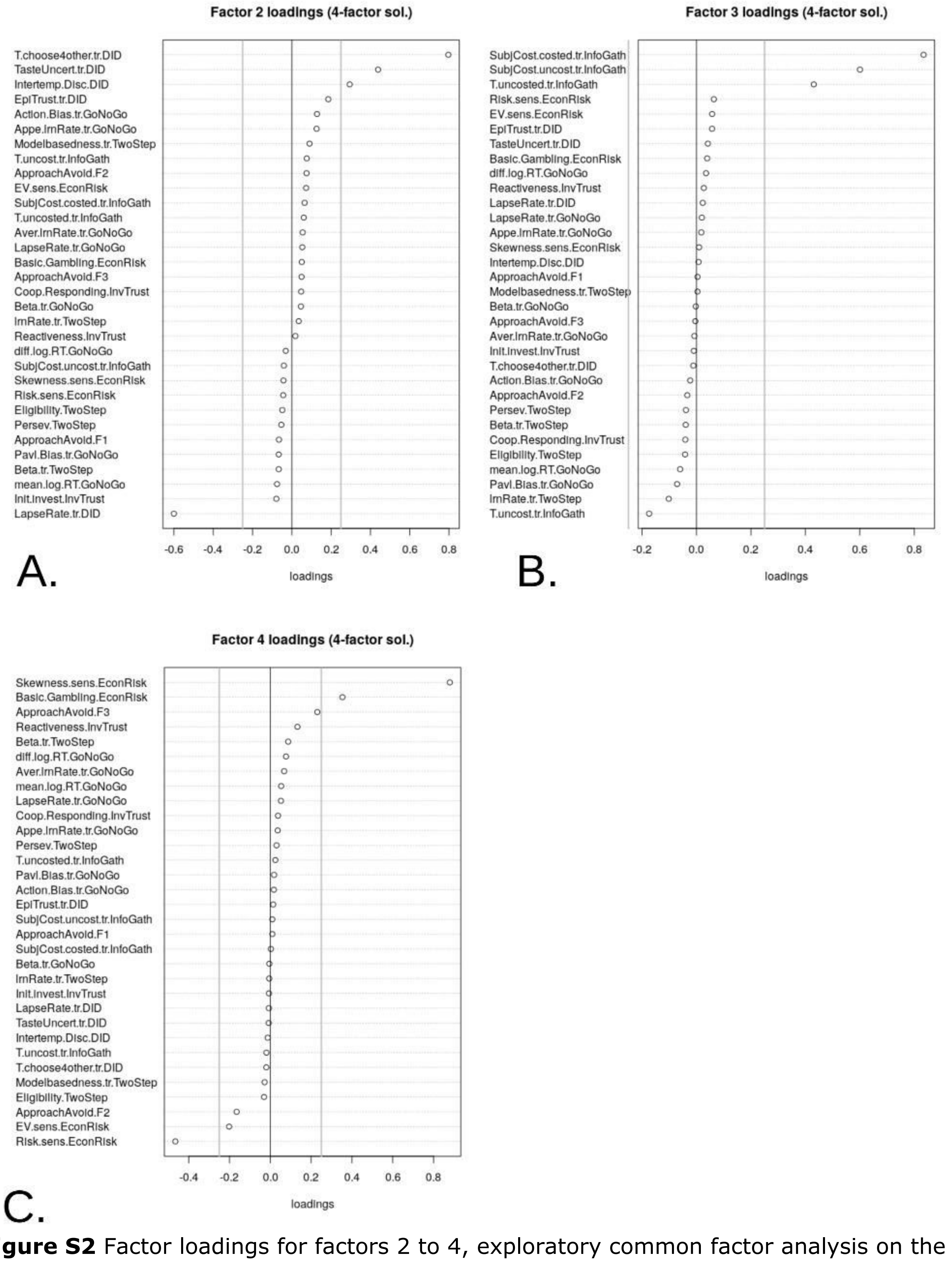


**Figure S2** Factor loadings for factors 2 to 4, exploratory common factor analysis on the whole sample. loadings with absolute value over the noise floor or over 0.25 (gray lines) are exclusively from **A.** Delegated Discounting task for Factor 2 **B.** Information Gathering task for Factor 3 and **C.** Economic risk preference task for Factor 4.

#### B2. Stability Analysis

We examined construct stability of decision acuity by correlating component *d* scores on half the sample with the same scores derived from the first ECFA component on the other half of the sample.

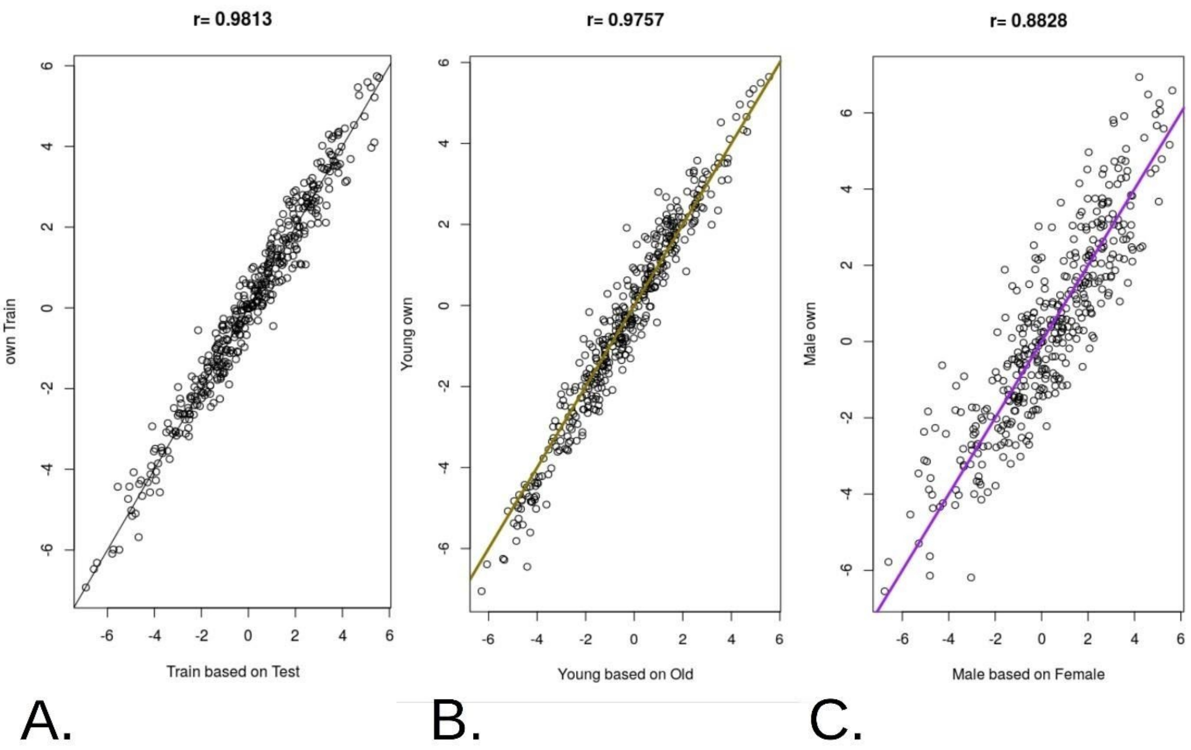


**Figure S3** Stability of the construct of decision acuity with respect to random variation in the data, age or sex. In each case, component factor scores for half the sample based on ECFA of that same half-sample is predicted by component scores for the same individuals, but based on the construct (i.e., factor loadings) derived from the opposite half of the data. **A.** Exploratory-confirmatory split gives a very high correlation ($r=0.98$, $p \cong 0.0$) attesting to the reliability of the construct **B.** Median split at age= 18.54 years. Very high correlation ($r=0.98$, $p \cong 0.0$) attests to the stability of the construct in young adults vs. teenagers. **C.** Female-male split shows somewhat lower correlation ($r=0.88$, $p \cong 0.0$), suggesting that the same 'average' construct can be used in both sexes, but also that subtle sexual dimorphism exists.

Here we were not primarily interested in the factor structure of decision-making, but in the stability of the construct of decision acuity. We thus divided the sample into two subgroups either by age (at 19) or by sex. We argued that if the construct itself was stable across age (and sex), then the decision acuity factor score for each participant could be calculated either using the factor loadings derived from the participant's own group or indeed the opposite one. Individuals with substantially differing scores would indicate that a different latent construct organized decision-making across the subgroups. If, for example, *d* was an invariant latent construct with respect to age, then the pattern of loadings derived from older participants would give the same scores when applied to younger participants as and ECFA on the young participant data themselves. *d* was highly stable across the discovery-confirmation random split (0.99 confidence interval for r(exploratory based on confirmatory loadings, own exploratory) = 0.976,0.985), as well as age CIr(young|old, own young)=0.969,0.9811). Its stability across gender was satisfactory but significantly lower, evidencing a small degree of sexual dimorphism CIr(male|female, own male)=0.820,0.887, Figure S3. Fit indicators were similar for the whole sample and for each split (e.g. RMSEA 90% CIs for females, males, younger, older and all were 0.051-0.061,0.052-0.062,0.054-0.064,0.046- 0.056 and 0.052-0.058 respectively)

As mentioned above, none of the analyses reported here was materially affected by excluding from the sample of 830 participants the 50 who had a diagnosis of DSM5 depression.

#### C. Additional associations of d with performance, symptoms and IQ

##### C1. *d*, performance and IQ

In order to check the interpretation that *d* reflects better decision-making, we first formed a performance measure across tasks. First, we excluded the discounting and Roulette tasks, as these specifically probed the balance of amounts won vs. other dimensions of the return, namely its delay and uncertainty respectively. Second, we excluded the Approach-Avoidance conflict task, as one of the measures by which it entered the estimation of *d* (ApproachAvoid.F3 in Figure 1 and Supplement Table S1 ) was judged to be too close to a performance measure already. Parenthetically, this measure loaded modestly in the expected direction onto *d*, i.e. positively, with a weight of 0.24 and high uniqueness (Figure 1). We then t-scored winnings within each of the Go-NoGo, Information Gathering, Investor-Trustee and Two-step tasks, and averaged these scores across tasks. The Pearson raw and partial correlation table between this task-performance measure, *d* and WASI total IQ had as follows, confirming the interpretation of *d* as conducive to profitable decision-making even above and beyond IQ.

| | | *d* | WASI IQ |
|---|---|---|---|
| task performance | partial | r=0.42, p <1e-10 | r=0.05, p=0.14 |
| | raw | r=0.50, p <1e-10 | r=0.30, p<1e-10 |
| *d* | partial | - | r=0.44, p<1e-10 |
| | raw | - | r=0.51, p<1e-10 |

**Table S2** Relations of d and IQ with overall performance in four key tasks.

##### C2. The association between *d* and symptoms is not explained by IQ

We tested whether each symptom factor was significantly associated with *d* while controlling for age and IQ raw scores, and found that in no case could the association of *d* with psychological scores be explained by IQ. First, 'Aberrant thinking', remained significantly associated with *d* (bz=-0.14, SE(bz)=0.045 , p=0.0018) after controlling for the IQ sub-scores. Here, vocabulary IQ also contributed (bz=-0.11, SE(bz)=0.044, p=0.016) but matrix IQ did not (p=0.96). Second, the association of *d* with 'Worry' remained significant (bz=+0.12, SE(bz)=0.044, p=0.0077) while in this regression matrix and vocabulary IQ were not (p=0.27 and 0.82 respectively). Third, 'Antisocial behaviour' remained significantly associated with *d,* (bz=-0.12, SE(bz)=0.043, p=0.0046), with matrix and vocabulary IQ not so (p=0.27 and 0.082).

### D. Brain connectivity analyses

#### D1. SPLS pipeline

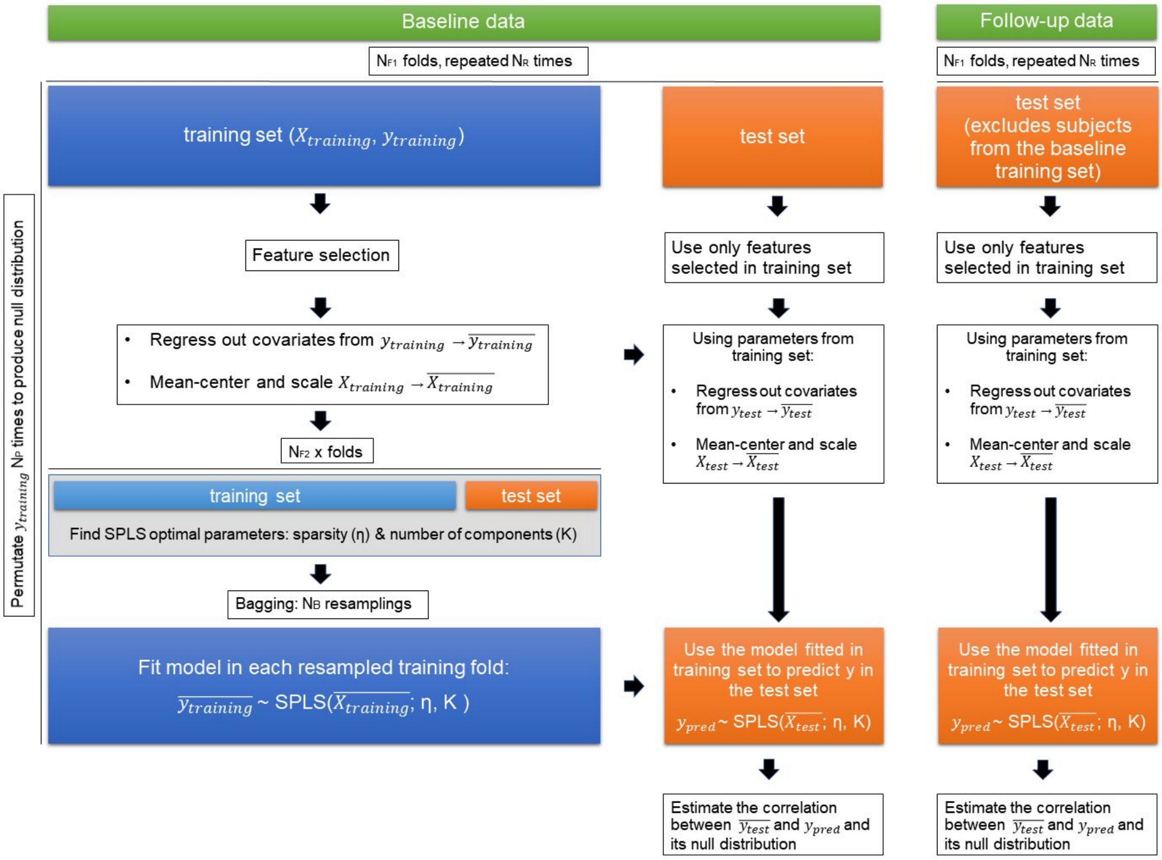


**Figure S4** Flow diagram of the nested cross-validation pipeline used to estimate how strongly decision acuity (similarly for IQ) could be predicted from brain data. Essentially, a predictive model was derived from training folds and then applied to the brain data from test folds to derive predicted values for the decision acuity for each individual. This could then be compared with the experimentally derived decision acuity. In our study, $N_B$ = 200, $N_{F1}$ = 20, $N_{F2}$ = 10, $N_R$ = 5, $N_P$ = 30, X corresponds to the rsFC features and y to the scores predicted (d or IQ).

### E - The Neuroscience in Psychiatry Consortium

#### Principal investigators:

Edward Bullmore (CI from 01/01/2017)
Raymond Dolan
Ian Goodyer (CI until 01/01/2017)
Peter Fonagy
Peter Jones

#### NSPN funded staff:

Michael Moutoussis
Tobias Hauser
Sharon Neufeld
Rafael Romero-Garcia
Michelle St Clair
Petra Vértes
Kirstie Whitaker
Becky Inkster
Gita Prabhu
Cinly Ooi
Umar Toseeb
Barry Widmer
Junaid Bhatti
Laura Villis
Ayesha Alrumaithi
Sarah Birt
Aislinn Bowler
Kalia Cleridou
Hina Dadabhoy
Emma Davies
Ashlyn Firkins
Sian Granville
Elizabeth Harding
Alexandra Hopkins
Daniel Isaacs
Janchai King
Danae Kokorikou
Christina Maurice
Cleo McIntosh
Jessica Memarzia
Harriet Mills
Ciara O'Donnell
Sara Pantaleone
Jenny Scott

#### Affiliated scientists:

Pasco Fearon

John Suckling
Anne-Laura van Harmelen
Rogier Kievit

### F - References to the Supplement


Bach, D. R., Guitart-Masip, M., Packard, P. A., Miró, J., Falip, M., Fuentemilla, L., & Dolan, R. J. (2014). Human hippocampus arbitrates approach-avoidance conflict. *Current Biology*, *24*(5), 541–547.

Bickel, W. K., Jarmolowicz, D. P., Mueller, T., Koffarnus, M. N., & Gatchalian, K. M. (2012). Excessive discounting of delayed reinforcers as a trans-disease process contributing to addiction and other disease-related vulnerabilities: Emerging evidence. *Pharmacology and Therapeutics*, *134*, 287–297. https://doi.org/10.1016/j.pharmthera.2012.02.004

Daw, N. D., Gershman, S. J., Seymour, B., Dayan, P., & Dolan, R. J. (2011). Model-based influences on humans' choices and striatal prediction errors. *Neuron*, *69*(6), 1204–1215.

Dayan, P. (2014). Rationalizable irrationalities of choice. *Topics in Cognitive Science*, *6*(2), 204–228.

Fett, A.-K. J., Shergill, S. S., Joyce, D. W., Riedl, A., Strobel, M., Gromann, P. M., & Krabbendam, L. (2012). To trust or not to trust: the dynamics of social interaction in psychosis. *Brain*, *135*(3), 976–984.

Guitart-Masip, M., Fuentemilla, L., Bach, D. R., Huys, Q. J. M., Dayan, P., Dolan, R. J., & Duzel, E. (2011). Action dominates valence in anticipatory representations in the human striatum and dopaminergic midbrain. *The Journal of Neuroscience*, *31*(21), 7867–7875.

Kerr, M., Van Zalk, M., & Stattin, H. (2012). Psychopathic traits moderate peer influence on adolescent delinquency. *J Child Psychol Psychiatry*, *53*, 826–835. https://doi.org/10.1111/j.1469-7610.2011.02492.x

King-Casas, B., Sharp, C., Lomax-Bream, L., Lohrenz, T., Fonagy, P., & Montague, P. R. (2008). The rupture and repair of cooperation in borderline personality disorder. *Science*, *321*(5890), 806–810. Retrieved from http://eutils.ncbi.nlm.nih.gov/entrez/eutils/elink.fcgi?cmd=prlinks&dbfrom=pubmed&retmode=ref&id=18687957

Lincoln, T. M., Peter, N., Schafer, M., & Moritz, S. (2010). From stress to paranoia: an experimental investigation of the moderating and mediating role of reasoning biases. *Psychol Med*, *40*(1), 169–171. Retrieved from http://eutils.ncbi.nlm.nih.gov/entrez/eutils/elink.fcgi?cmd=prlinks&dbfrom=pubmed&retmode=ref&id=19671213

Lincoln, T. M., Ziegler, M., Mehl, S., & Rief, W. (2010). The jumping to conclusions bias in delusions: specificity and changeability. *J Abnorm Psychol*, *119*(1), 40–49. Retrieved from http://eutils.ncbi.nlm.nih.gov/entrez/eutils/elink.fcgi?cmd=prlinks&dbfrom=pubmed&retmode=ref&id=20141241

Moutoussis, M., Bentall, R. P., El-Deredy, W., & Dayan, P. (2011). Bayesian modeling of Jumping-to-Conclusions Bias in delusional patients. *Cognitive Neuropsychiatry*.

Moutoussis, M., Dolan, R. J., & Dayan, P. (2016). How people use social information to find out what to want in the paradigmatic case of inter-temporal preferences. *PLOS Computational Biology*.

Nicolle, A., Klein-Flugge, M., Hunt, L. T., Vlaev, I., Dolan, R. J., & Behrens, T. E. J. (2012). An Agent Independent Axis for Executed and Modeled Choice in Medial Prefrontal Cortex. *Neuron*, *75*, 1114–1121.

Sripada, C. S., Angstadt, M., Banks, S., Nathan, P. J., Liberzon, I., & Phan, K. L. (2009). Functional neuroimaging of mentalizing during the trust game in social anxiety disorder. *Neuroreport*, *20*(11), 984–989. Retrieved from http://eutils.ncbi.nlm.nih.gov/entrez/eutils/elink.fcgi?cmd=prlinks&dbfrom=pubmed&retmode=ref&id=19521264

Symmonds, M., Wright, N. D., Bach, D. R., & Dolan, R. J. (2011). Deconstructing risk: separable encoding of variance and skewness in the brain. *Neuroimage*, *58*(4), 1139–1149.
